# A new transgenic reporter line reveals Wnt-dependent Snail2 reexpression and cranial neural crest differentiation in *Xenopus*

**DOI:** 10.1101/520726

**Authors:** Jiejing Li, Mark Perfetto, Christopher Materna, Rebecca Li, Hong Thi Tran, Kris Vleminckx, Melinda K. Duncan, Shuo Wei

**Affiliations:** Department of Biology, West Virginia University, Morgantown, WV 26506, USA; Department of Biological Sciences, University of Delaware, Newark, DE 19716, USA; Brown University, Providence, RI 02912, USA; Department for Molecular Biomedical Research and Center for Medical Genetics, Ghent University, B-9052 Ghent, Belgium; Department of Clinical Laboratory, The Affiliated Hospital of KMUST, Medical School, Kunming University of Science and Technology, Kunming 650032, China

## Abstract

During vertebrate embryogenesis, the cranial neural crest (CNC) forms at the neural plate border and subsequently migrates and differentiates into many types of cells. The transcription factor Snail2, which is induced by canonical Wnt signaling to be expressed in the early CNC, is pivotal for CNC induction and migration in *Xenopus*. However, *snail2* expression is silenced during CNC migration, and its roles at later developmental stages remain unclear. We generated a transgenic *X. tropicalis* line that expresses enhanced green fluorescent protein (eGFP) driven by the *snail2* promoter/enhancer, and observed eGFP expression not only in the pre-migratory and migrating CNC, but also the differentiating CNC. This transgenic line can be used directly to detect deficiencies in CNC development at various stages, including subtle perturbation of CNC differentiation. In situ hybridization and immunohistochemistry confirm that Snail2 is reexpressed in the differentiating CNC. Using a separate transgenic Wnt reporter line, we show that canonical Wnt signaling is also active in the differentiating CNC. Blocking Wnt signaling shortly after CNC migration causes reduced *snail2* expression and impaired differentiation of CNC-derived head cartilage structures. These results suggest that Wnt signaling drives the reexpression of *snail2* in the post-migratory CNC and regulates CNC differentiation.

## Introduction

The cranial neural crest (CNC) cells are a transient group of multipotent stem cells that exists during early vertebrate embryogenesis. CNC development can be divided into three major stages: the induction, migration and post-migratory differentiation of the CNC. During gastrulation, the CNC is induced at the posterior neural plate border (NPB) between the neuroectoderm and the epidermis. The CNC cells continue to proliferate and undergo epithelial-mesenchymal transition (EMT), and subsequently emigrate from the closing neural tube in several streams that target distinct areas. Once the migrating CNC cells arrive at their destinations, they begin to differentiate into multiple types of cells that contribute to various tissues. Derivatives of CNC include nearly all the craniofacial structures, such as skeleton, connective tissues, muscles and the peripheral nervous system^1–3^.

The ability to differentiate into multiple cell types and contribute to many tissues makes CNC an intriguing subject of research in developmental biology. Recent studies have uncovered several new types of cells that may derive from the CNC^4,5^, but the identification of all CNC derivatives remains a daunting task and requires new lineage-tracing tools. Moreover, CNC development is a dynamic and complex process that is tightly controlled spatially and temporally. Perturbation of CNC cells during any developmental stages may result in defects known as neurocristopathies, which are among the most common birth defects in humans^6^. Some of these defects are subtle, and are therefore difficult to detect with the techniques currently available. Hence the development of model systems that can be used for tracing CNC development will have a tremendous impact on the studies of CNC biology and the etiology of neurocristophathies. To this end, a number of transgenic mouse lines have been generated to facilitate tracing of the CNC lineage^7^. However, due to technical difficulties and possibly higher levels of functional redundancy in mice, non-mammalian vertebrates are often the preferred models for studying CNC development^8^. Among the non-mammalian models, zebrafish and *Xenopus* are particularly suitable for live imaging of tissue morphogenesis, owing to their external embryonic development, transparent epithelium and large brood size. Transgenic zebrafish lines expressing eGFP or Cre recombinase driven by the *sox10* promoter have been widely used as lineage tracing tools to label neural crest derivatives^9,10^. Historically, the detection of CNC in *Xenopus* was almost solely dependent on in situ hybridization for CNC markers such as *snail2, sox9* and *twist*, whereas Alcian blue staining was commonly used for visualization of the head cartilage structures that derive from the CNC. These procedures are time-consuming and labor-intensive, and require fixation that prevents further manipulations of the embryos. While this manuscript was being prepared, Alkobtawi et al. published the generation of *pax3-GFP* and *sox10-GFP*, the first two *X. laevis* transgenic lines that can be used for live imaging of CNC induction and migration, respectively^11^. However, currently no similar tool is available for *X. tropicalis*, a diploid species that is highly suitable for genetic studies, or for imaging CNC differentiation in any frog species.

Snail2 (a.k.a. Slug) is a zinc-finger transcription factor that is expressed in early CNC precursors and is required for the induction/specification of CNC in Xenopus^12,13^. A major signaling pathway that activates *snail2* expression during CNC induction is the canonical Wnt (hereinafter referred to as “Wnt”) pathway, as forced activation of Wnt signaling causes ectopic expression of *snail2* and other CNC markers, and blocking Wnt signaling inhibits *snail2* expression and CNC induction^14^. Importantly, the *snail2* enhancer contains an evolutionarily conserved LEF-TCF binding site and can respond to Wnt signaling, suggesting that *snail2* is a direct Wnt target gene^15^. After CNC induction, *snail2* continues to be expressed in the pre-migratory and early migrating CNC, and plays a critical role in EMT and migration of the CNC^13,16^. However, recent quantitative RT-PCR and RNA-seq data show that *snail2* expression drastically decreases in *Xenopus* embryos after the CNC cells begin to migrate^17–19^. Therefore, studies published to date have been focused on the roles of Snail2 in CNC induction and migration, and little is known about the expression or function of this important transcription factor at later stages of embryonic development.

Because of its specific expression and pivotal function during both CNC induction and migration, *snail2* is one of the most commonly used CNC markers in frogs and other vertebrates such as chicks. The cis-regulatory elements of *X. tropicalis snail2* gene have been well characterized, and a ~3.9 kb region of the promoter/enhancer sequence has been shown to contain the LEF/TCF-binding site and be able to drive CNC-specific GFP expression when transiently expressed^15^. Using the I-SceI meganuclease-mediated transgenic method^20^, we generated a transgenic line that expresses eGFP driven by this ~3.9 kb *snail2* promoter/enhancer. Expression of eGFP in the *snail2::egfp* transgenic embryos not only faithfully reflects the expression of endogenous *snail2* in the pre-migratory and early migrating CNC, but also unveils a previously unknown expression of *snail2* in the post-migratory CNC. In the *snail2::egfp* transgenic tadpoles, eGFP labels multiple differentiating CNC derivatives, and subtle perturbation of CNC differentiation, such as those caused by partial knockdown of the disintegrin metalloproteinase ADAM13, can be readily detected using the *snail2::egfp* transgenic embryos. We further show that Wnt signaling, which is regulated by ADAM13, is similarly activated in the differentiating CNC. Finally, blocking Wnt signaling using small-molecule inhibitors shortly after the completion of CNC migration leads to reduction in *snail2* expression and under-differentiation of CNC-derived head cartilage structures, suggesting that Wnt is required for post-migratory CNC differentiation, probably by regulating *snail2* expression.

## Results

### Generation of the *snail2::egfp* transgenic *X. tropicalis* line

We cloned a ~3.9-kb *X. tropicalis* genomic DNA upstream of the *snail2* transcription start site, including the promoter and the 5’-enhancer, as described by Vallin et al.^15^. This *snail2* promoter/enhancer sequence was inserted into a transgenic vector (Fig. S1A), and stable transgenic founders were generated as described by Ogino et al.^20^. At stage ~22, some of these founder embryos showed distinct fluorescence in the migrating CNC with minimal ectopic expression (Fig. S1B). This is consistent with *snail2* expression in the migrating CNC, and suggests that the reporter construct had been integrated into the genome in these embryos^20,21^. When a potential transgenic founder was crossed with wild-type frogs, it produced heterozygous progeny (F1) that showed distinct fluorescence patterns (see below), indicating that the transgene insertion was inherited through germline transmission. We further inbred the F1 transgenic frogs to produce F2 progeny. About 71% (161/226) of all F2 embryos were eGFP-positive, and ~25% (41/161) of eGFP-positive embryos displayed stronger fluorescence than the others. The embryos with higher eGFP expression were singled out and raised to sexual maturity, and further crossing with wild-type frogs yielded 100% eGFP-positive embryos, suggesting that these frogs with higher eGFP expression were homozygotes. These results point to a single integration of the transgene, which was confirmed by whole-genome sequencing (see below). All heterozygous and homozygous *snail2::egfp* transgenic frogs were healthy and fertile, and displayed normal craniofacial morphology (data not shown). In situ hybridization for *snail2* and *sox9* in the pre-migratory CNC (Fig. S2A, B, D), as well as *snail2* and *twist* in the migrating CNC (Fig. S2C, E, F), also showed normal patterns, indicating that the transgene insertion did not affect CNC development. To better understand the potential impact of the transgene insertion at the molecular level, we carried out whole-genome shotgun sequencing on heterozygous *snail2::egfp* transgenic embryos, and mapped the transgene insertion to a single non-coding region on Chromosome 1 (Wang and Wei, manuscript in preparation).

### eGFP is expressed in the CNC lineage including differentiating CNC in *snail2::egfp* transgenic embryos

The *snail2::egfp* transgenic embryos displayed highly specific fluorescence patterns. Green fluorescence was observed in the pre-migratory CNC cells as early as stage ~15 (Fig. 1A), and in the migrating CNC streams at stage 19-22 (Fig. 1B, C). Although recent reports show that *snail2* expression is downregulated during CNC migration in *Xenopus* embryos^17–19^, fluorescence signals were clearly detectable in the post-migratory CNC as well as the developing eyes and brain at tailbud stages in the *snail2::egfp* transgenic embryos (Fig. 1D, D’). These patterns continued into swimming tadpole stages, when eGFP labeled multiple structures that are known to have CNC contributions (Fig. 1E-H’). In particular, the morphological changes were highlighted by green fluorescence in the *snail2::egfp* embryos as cells in the pharyngeal arches developed from condensing mesenchyme (Fig. 1D) to partially differentiated cartilage (Fig. 1E’, F’, G’), and eventually to highly complex cartilaginous structures (Fig. 1H’). Starting from stage ~42, fluorescence was also detected in several cranial nerves, including the trigeminal and oculomotor nerves, as well as the olfactory nerves that are known to have a CNC contribution in other vertebrate species (Fig. 1E’, F’, G, G’, H’)^4^. Recent lineage tracing studies suggest that CNC may also contribute to some of the neurons in the olfactory epithelium in zebrafish and mice^22,23^. At stage ~42, fluorescence became detectable in the emerging olfactory epithelium of the *snail2::egfp* tadpoles (Fig. 1E, E”), and the signal increased throughout early tadpole stages and persisted to later stages (Fig. 1F, G, H). We also observed strong fluorescence in the developing thymus, which is known to be populated by CNC cells^24^, at stage ~53 (Fig. 1H). Although *snail2* was not shown previously to be expressed in the post-migratory CNC, the eGFP signals likely reflect the true expression of *snail2* instead of simply being the remnant of early expression prior to silencing during CNC migration, as these eGFP signals remained in the differentiating CNC lineage for more than 2 weeks, whereas the half-life of eGFP is merely ~26 hr^25^. Additionally, fluorescence was found in the eyes and somites of the transgenic tadpoles (Fig. 1E’, F”, G’, H), consistent with published results showing that *snail2* is expressed in these tissues in various vertebrate species, ranging from fish to mice^26,27^.

**Figure 1.**
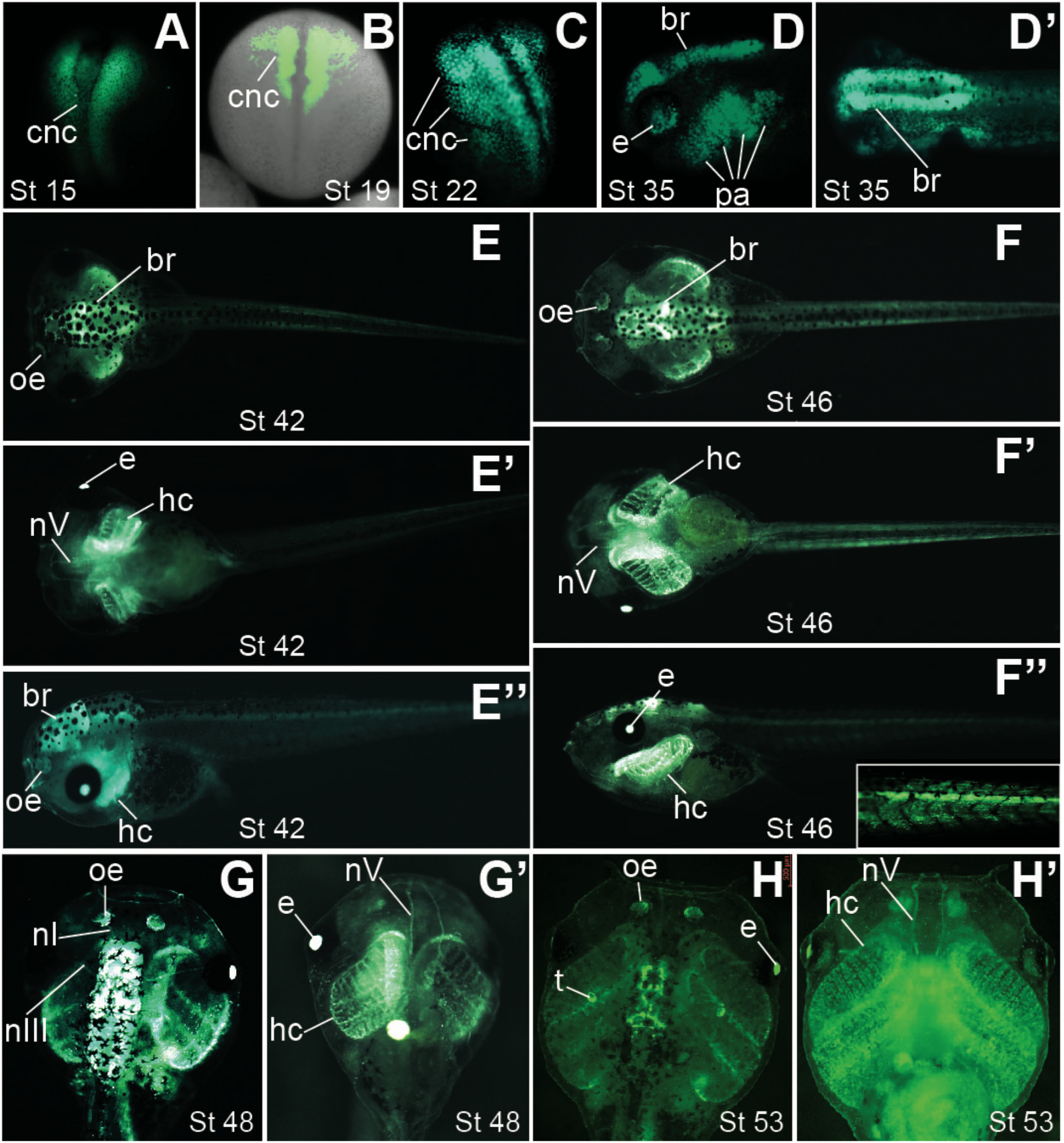
The *snail2::egfp* transgenic embryos show eGFP expression in the CNC lineage at various stages. Heterozygous *snail2::egfp* embryos were imaged at the indicated stages. **A-C**. Neurula-stage embryos showing eGFP expression in pre-migratory (**A**), early migrating (**B**) and extensively migrating (**C**) CNC. Green fluorescence and bright-field images are merged in B to show the relative positions of migrating CNC streams in the whole embryo. **D** and **D’**. Side (**D**) and dorsal (**D’**) views of a stage ~35 tadpole with eGFP expression in the brain (br), eye (e), and CNC cells forming condensing mesenchyme in the pharyngeal arches (pa). **E-E”**. Dorsal (**E**), ventral (**E’**) and side (**E”**) views of a stage ~42 tadpole. eGFP expression is detectable in the developing olfactory epithelium (oe), trigeminal nerves (nV), and CNC cells in the pharyngeal arches that begin to differentiate into head cartilage (hc). **F-F”**. Dorsal (**F**), ventral (**F’**) and side (**F”**) views of a stage ~46 tadpole. eGFP is seen in the more differentiated trigeminal nerves and head cartilage structures. Inset in **F”** is a higher-magnification image showing eGFP expression in tail somites. **G** and **G’**. Dorsal (**G**) and ventral (**G’**) views of a stage ~48 tadpole, with eGFP visible in both the olfactory (nI) and oculomotor (nIII) nerves. **H** and **H’**. Dorsal (**H**) and ventral (**H’**) views of a stage ~53 tadpole. eGFP labels the thymus (t) and highly differentiated head cartilage structures.

### eGFP patterns in the *snail2::egfp* transgenic embryos faithfully reflect the endogenous expression of Snail2 mRNA and protein

Because the stability of eGFP prevents the observation of subtle dynamics of *snail2* expression in the migrating and post-migratory CNC, we carried out in situ hybridization to detect endogenous *snail2* transcripts at various developmental stages. At stage ~12, *snail2* mRNA was mainly expressed in the midline; there was also weak expression in the future CNC territory (Fig. 2A). About half an hour later (stage ~12.5), when the embryos approached the end of gastrulation, strong *snail2* expression was detected in the newly formed CNC (Fig. 2B). At early neurula stages, *snail2* continued to be expressed in the pre-migratory CNC, but the midline expression diminished (Fig. 2C). By stage ~19, CNC cells had emigrated from the closing neural tube, as shown by the in situ hybridization for *snail2* (Fig. 2D-F). Therefore, the patterns of eGFP in the *snail2::egfp* transgenic embryos faithfully reflect the expression of *snail2* in both pre-migratory and early migrating CNC. The expression of *snail2* persisted in the early migrating CNC cells, which formed distinct streams as they migrated out of the completely closed neural tube, but the intensity started to decrease thereafter and was minimal at stage ~31, several stages after CNC cells ceased migration (Fig. 2F, G and data not shown). This is in line with previous reports that *snail2* expression is downregulated in late migrating CNC cells^17–19^. However, *snail2* expression started to increase again at stage ~32 and was clearly detectable in the eyes as well as the condensing mesenchyme within the pharyngeal arches (Fig. 2H). At swimming tadpole stages, *snail2* transcripts were found in the head and brain as well as the somites (Fig. 2I, J). Unfortunately, visualization of detailed cartilage structures was difficult, probably because the in situ probes and/or alkaline phosphatase substrate were trapped in the cavities that had formed in the head at these stages. We therefore dissected the head cartilage, and were able to detect *snail2* staining in the fine cartilaginous structures (insets in Fig. 2I, J), which was highly similar to the eGFP patterns in the *snail2::egfp* transgenic tadpoles (Fig. 1E’, F’). Thus, *snail2* is re-expressed in the differentiating CNC cells after migration.

**Figure 2.**
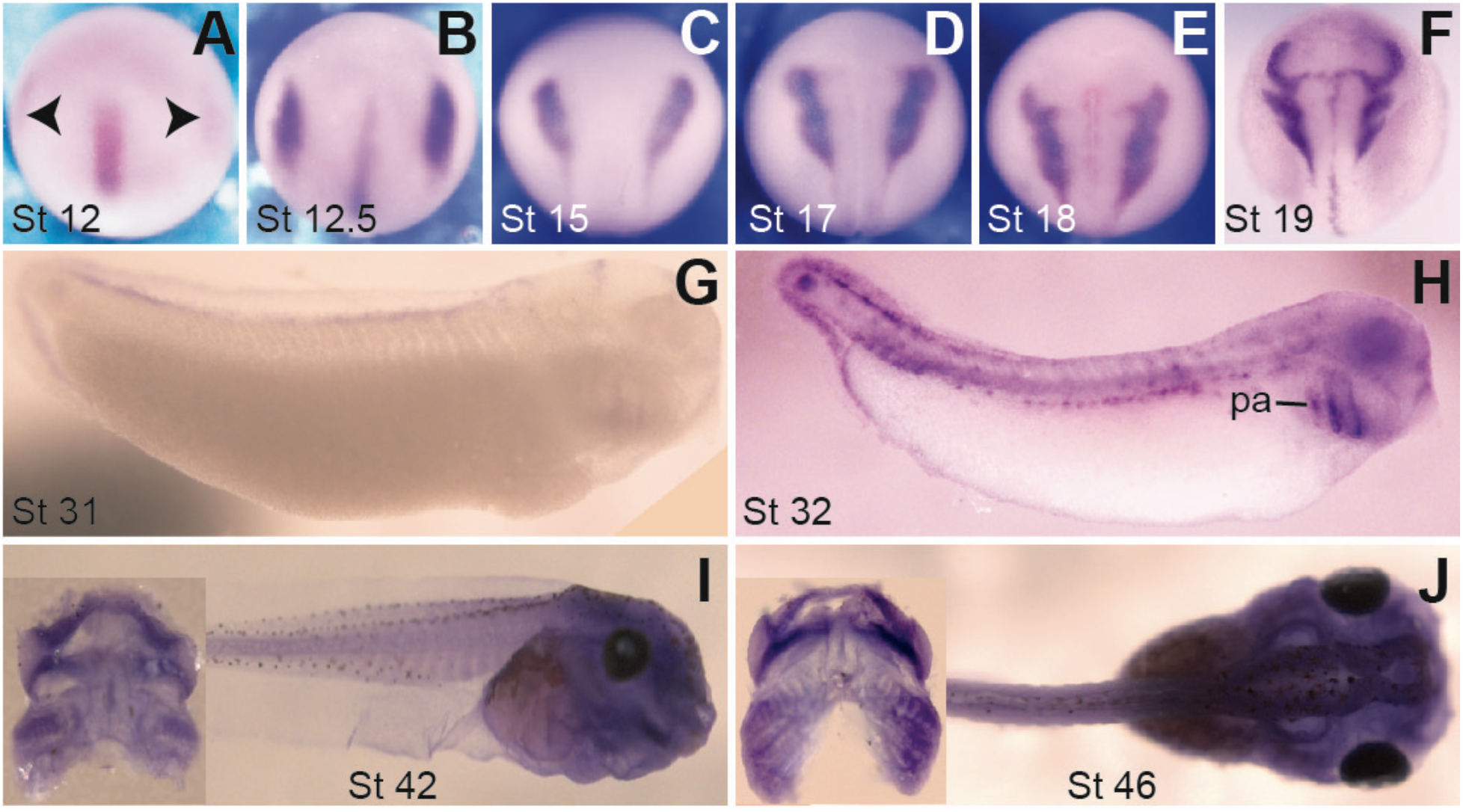
The transcripts of *snail2* are expressed in pre-migratory, migrating and post-migratory CNC. In situ hybridization was performed for *snail2* with wild-type *X. tropicalis* embryos at the indicated stages. Arrowheads in **A** indicate weak *snail2* expression in the emerging CNC. *Snail2* continues to be expressed in the pre-migratory (**B** and **C**) as well as migrating (**D-F**) CNC cells. The expression is silenced by stage ~31 (**G**), but restarts in the CNC cells that form condensing mesenchyme in the pharyngeal arches (pa; **H**) and persists in the differentiating head cartilage structures (**I** and **J**). Insets in **I** and **J** are head cartilage dissected from tadpoles after in situ hybridization.

We further performed double-immunohistochemistry to determine if the spatiotemporal patterns of eGFP in the *snail2::egfp* embryos reflect those of the endogenous Snail2 protein. At stage ~17, eGFP and Snail2 displayed significant co-localization in the closing neural tube, where some CNC cells just started to emigrate (Fig. 3A-A”). At stage ~46, eGFP clearly co-localized with Snail2 in the differentiating head cartilage structures, trigeminal nerves, olfactory epithelium, eye and brain (Fig. 3B-C”). It should be noted, though, that there are regions where eGFP but not Snail2 was detected, which may reflect the difference in stability between these two proteins. Together with the in situ hybridization data, these results indicate that Snail2 mRNA and protein are expressed in the post-migratory CNC, and that the *snail2::egfp* transgenic line is suitable for tracing the CNC lineage at various developmental stages.

**Figure 3.**
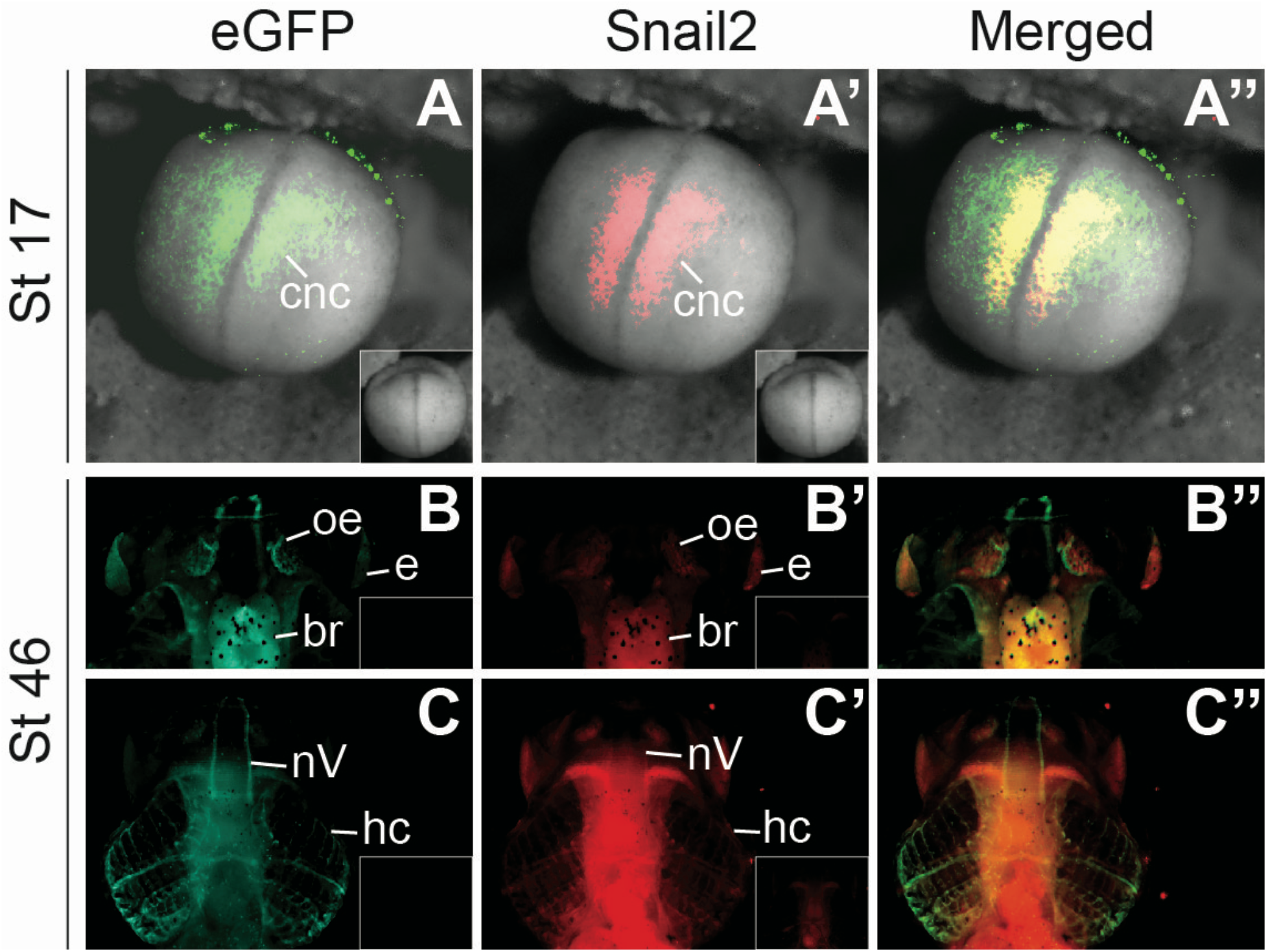
Co-localization of eGFP and Snail2 proteins in *snail2::egfp* embryos. Immunohistochemistry was carried out for eGFP (green) and Snail2 (red) simultaneously at the indicated stages in *snail2::egfp* embryos and tadpoles. **A-A”**. eGFP and Snail2 are co-localized in the CNC at the onset of migration. A control embryo processed with secondary antibodies only but not either primary antibody did not display any signal (insets in A and A’). **B-C”**. Dorsal (**B-B”**) and ventral (**C-C”**) views of a stage ~46 tadpole showing co-localization of eGFP and Snail2 in the head cartilage (hc), brain (br), eye (e), trigeminal nerve (nV), and olfactory epithelium (oe).

### CNC defects at various stages are readily detectable in live *snail2::egfp* transgenic embryos

We next tested if the *snail2::egfp* line can be used for real-time detection of CNC defects. To do this we carried out antisense morpholino (MO)-mediated knockdown of the disintegrin metalloproteinase ADAM13, a protease that is known to be required for normal CNC induction and migration^28–30^. An ADAM13 MO (MO 13-3), which has been well characterized in previous studies^28,30,31^, was injected into one blastomere of 8-cell stage *snail2::egfp* embryos to target the dorsal-animal region; a red-fluorescence dye was co-injected as lineage tracer. As expected, many embryos that were injected with ADAM13 MO (the “morphants”) show reduced eGFP expression on the injected side prior to CNC migration (Fig. 4A), suggesting that CNC induction was inhibited by ADAM13 knockdown. At stage ~22, CNC migration was also blocked in most embryos, as shown by the lack of eGFP labeled cells that emigrate from the neural tube in some ADAM13 morphants (Fig. 4B). To investigate if ADAM13 also affects post-migratory CNC development, we selected ADAM13 morphants with no apparent defects in CNC induction or migration, and cultured them to stage ~46. At this stage, various defects in CNC derivatives, such as reduction of the head cartilage structures and/or cranial nerves, were observed in some ADAM13 morphants (Fig. 4C-F). Interestingly, some of these morphants displayed hypoplasia or impaired differentiation of specific tissues that may have CNC contribution. For example, in a morphant with intact head cartilage structures and cranial nerves, we found that the olfactory epithelium was almost completely missing on the injected side (Fig. 4C, C’), suggesting that post-migratory CNC development, but not earlier CNC induction or migration, might be affected. Hence the *snail2::egfp* transgenic embryos can be used for live imaging of both early and late defects in CNC development.

**Figure 4.**
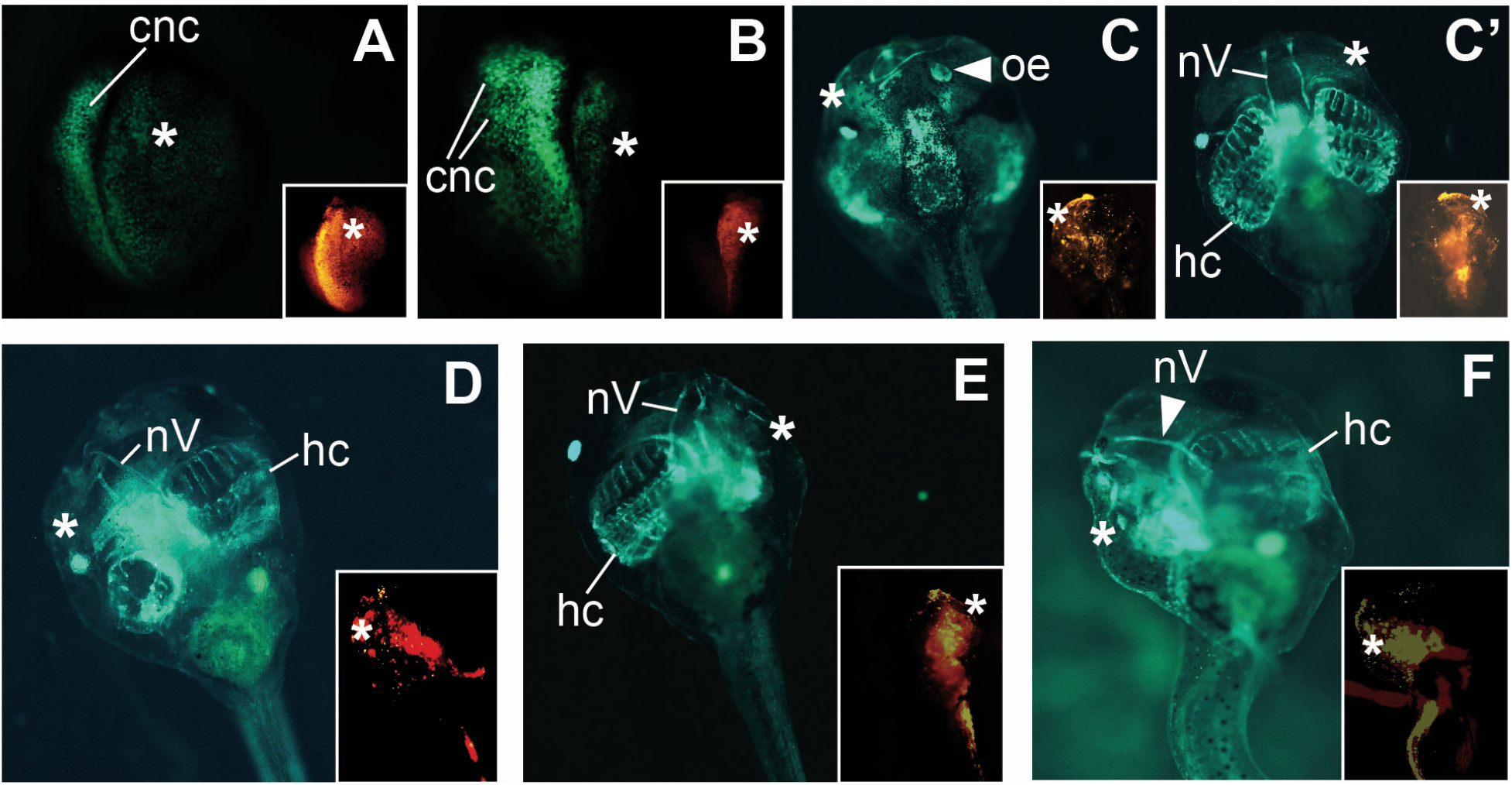
Phenotypes of ADAM13 knockdown displayed by *snail2::egfp* embryos. Eight-cell stage heterozygous *snail2::egfp* embryos were injected with 1.5 ng MO 13-3 to target ADAM13 in one dorsal-animal blastomere, and cultured to the indicated stages; a red fluorescent dye was co-injected as a lineage tracer. The injected side is denoted with a white asterisk, and structures that are present on the uninjected side but absent on the injected side are denoted with white arrowheads. Insets show red fluorescence images of the same embryos. **A** and **B**. Stage ~18 (**A**) and ~22 (**B**) embryos displaying reduced CNC domain on the injected side, as determined by eGFP expression. In B, CNC migration is normal on the uninjected side but inhibited on the injected side. **C-F**. Injected embryos that did not show apparent defects in CNC induction or migration were selected and cultured to stage ~46. **C** and **C’** are dorsal and ventral views, respectively, of the same tadpole. The olfactory epithelium (oe) is not detectable (**C**) but head cartilage (hc) and trigeminal nerve (nV) appear normal (**C’**) on the injected side of this embryo. **D** and **E**. Embryos with under-differentiated head cartilage structures on the injected side, as compared with uninjected side. **F**. An embryo with severely defective trigeminal nerve and head cartilage structures on the injected side.

### Wnt signaling is active in the post-migratory CNC

During CNC induction, ADAM13 functions by regulating Wnt signaling and *snail2* expression ^28,30^. *Snail2* is thought to be a direct Wnt target gene at this early stage of CNC development, because its 5’-enhancer contains a LEF/TCF binding site that can respond to Wnt signaling^1,15^. This LEF/TCF binding site was part of the 3.9 kb promoter-enhancer sequence that we used to generate the *snail2::egfp* transgenic line, raising the possibility that the re-expression of *snail2* in the post-migratory CNC, as reflected by eGFP patterns in the *snail2::egfp* embryos, is also induced by Wnt signaling. The effects of ADAM13 knockdown on post-migratory CNC development (Fig. 4C-G) further suggest that Wnt may have an important role in this later developmental process. To investigate the activity of Wnt signaling in the CNC at various developmental stages, especially during post-migratory CNC differentiation, we used a transgenic *X. tropicalis* Wnt reporter line that expresses destabilized eGFP driven by an artificial enhancer containing 7 LEF/TCF binding sites. This destabilized eGFP molecule has a short halflife (~2 hr) and can precisely reflect the dynamic on-and-off patterns of endogenous Wnt activity^32,33^. As shown previously, there was a strong Wnt signal at the posterior NPB during CNC specification (stage ~12.5; Fig. 5A), and this signal is critical for inducing *snail2* expression and the specification of CNC lineage^30,33^. This Wnt signal remained in the pre-migratory CNC, and was evident in the migrating CNC streams (Fig. 5B, C). Interestingly, strong Wnt signal was detected in the condensing mesenchyme in the pharyngeal arches at stage ~35, when the CNC cells start to differentiate (Fig. 5D). At stage ~42, Wnt was active in the differentiating head cartilage structures, cranial nerves, olfactory epithelium, eye and brain (Fig. 5E, E’), consistent with previously reported activities and/or functions of Wnt signaling in these tissues in mice^34–37^. These patterns are also strikingly similar to those of eGFP in the *snail2::egfp* embryos as well as endogenous Snail2 mRNA and protein, suggesting a possible role for Wnt signaling in inducing *snail2* expression, not only in the pre-migratory and migrating CNC but also in the differentiating CNC, in *X. tropicalis* embryos.

**Figure 5.**
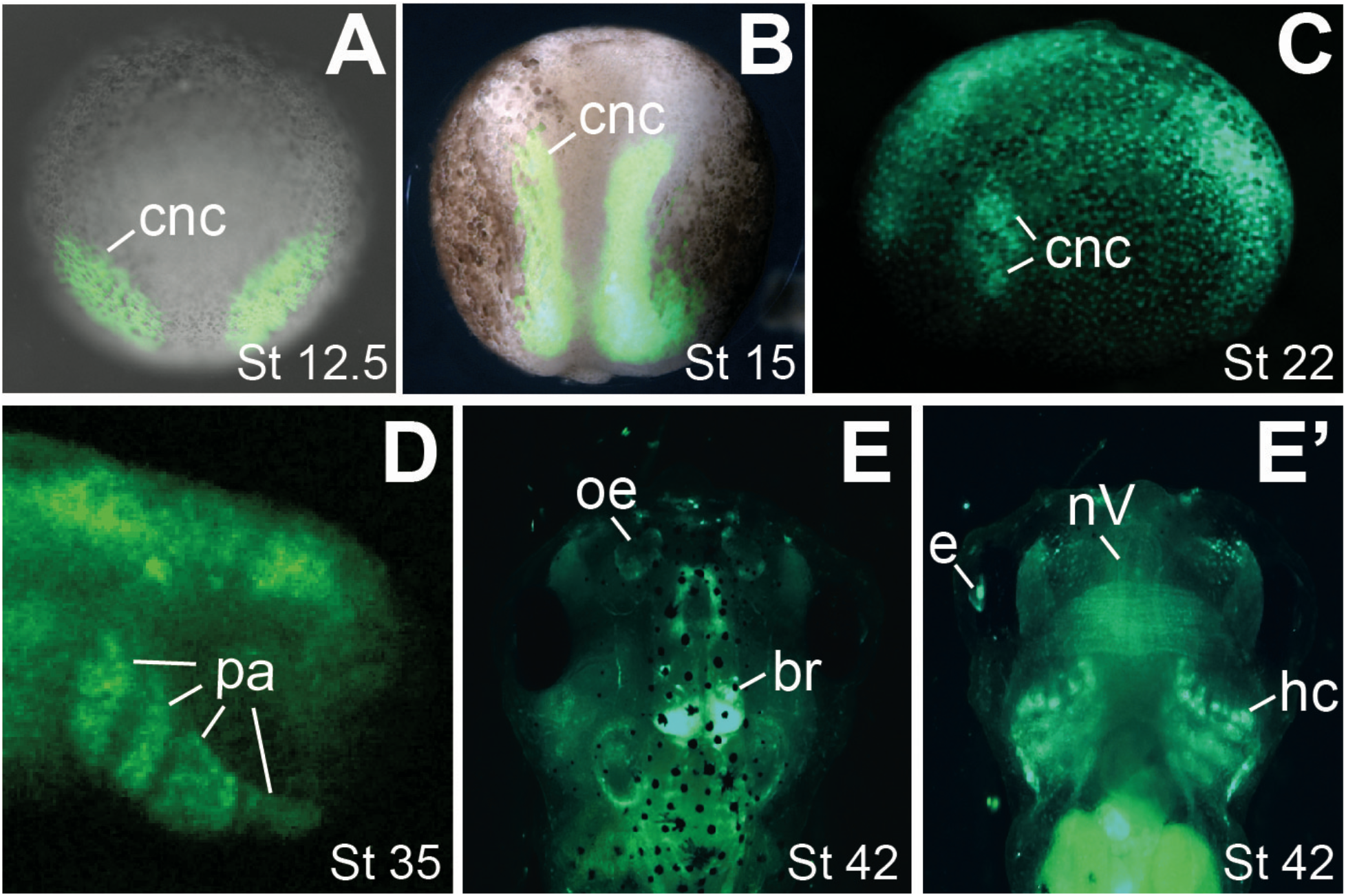
Wnt signaling activity in the CNC lineage. Heterozygous transgenic Wnt reporter embryos were imaged at the indicated stages. Expression of eGFP is detectable in the pre-migratory (**A** and **B**), migrating (**C**) and differentiating (**D-E’**) CNC. E and E’ are dorsal and ventral views, respectively, of the same tadpole. Green fluorescence and bright-field images are merged in **A** and **B** to show the relative positions of CNC in the whole embryo. br, brain; e, eye; hc, head cartilage; nV, trigeminal nerve; oe, olfactory epithelium; pa, pharyngeal arches.

### Wnt is required for head cartilage differentiation as well as *snail2* and *sox9* expression in the post-migratory CNC

We show above that some ADAM13 morphants displayed defects in the CNC lineage after CNC migration (Fig. 4C-F). Because ADAM13 is required for Wnt signal activation during CNC induction^28,30^, these results suggest that Wnt may be important for CNC differentiation. To further test this hypothesis, we treated transgenic embryos with small-molecule Wnt inhibitors starting at stage ~28, shortly after CNC completed migration. The treatment lasted until stage ~35, before CNC cells in the pharyngeal arches start to differentiate into cartilaginous structures (see Fig. 1D). Two Wnt inhibitors, which were identified in two independent screens, were used in these experiments. XAV939 stabilizes Axin, a major component of the β-catenin destruction complex, by inhibiting the tankyrases that stimulate Axin degradation^38^. The other compound, IWR1-endo, also elevates the protein levels of Axin, but the underlying mechanism remains unclear^39^. When *snail2::egfp* embryos were treated with high dosage of XAV939 or IWR1-endo after CNC migration, the branchial cartilage was clearly under-differentiated at stage ~44, as compared with embryos that were treated with vehicle control (Fig. 6A-D). In contrast, eGFP expression in the brain appeared to be normal (Fig. 6A’-C’). Similarly, defects in head cartilage structures were observed in Wnt reporter embryos treated with either Wnt inhibitor (Fig. 6E-H). Notably, global expression of the destabilized eGFP was greatly downregulated in these Wnt reporter embryos (Fig. 6E-G’), confirming that endogenous Wnt signaling was inhibited. Both the *snail2::egfp* and Wnt reporter tadpoles treated with Wnt inhibitors developed edema and died shortly after stage ~44. To rule out the possibility that the effects on head cartilage differentiation were secondary to the edema, we carried out similar treatment of the *snail2::egfp* embryos with lower dosage of Wnt inhibitors. These tadpoles did not have edema and were alive and swimming at stage ~47. However, the head cartilage structures were still underdeveloped as compared with those of the control embryos (Fig. 6I-L). No apparent reduction in total eGFP intensity was detected in the head cartilage of the *snail2::egfp* tadpoles at either stage ~44 or ~47 (Fig. 6A-C, I-K), suggesting that cell proliferation and death were not affected. The high eGFP levels seen in the *snail2::egfp* tadpoles upon Wnt inhibitor treatment is likely due to the stability of the eGFP protein, as endogenous *snail2* was found to be downregulated (see below). These results, together with the late CNC phenotypes displayed by the ADAM13 morphants (Fig. 4C-G), indicate that Wnt signaling plays critical roles in post-migratory CNC differentiation.

**Figure 6.**
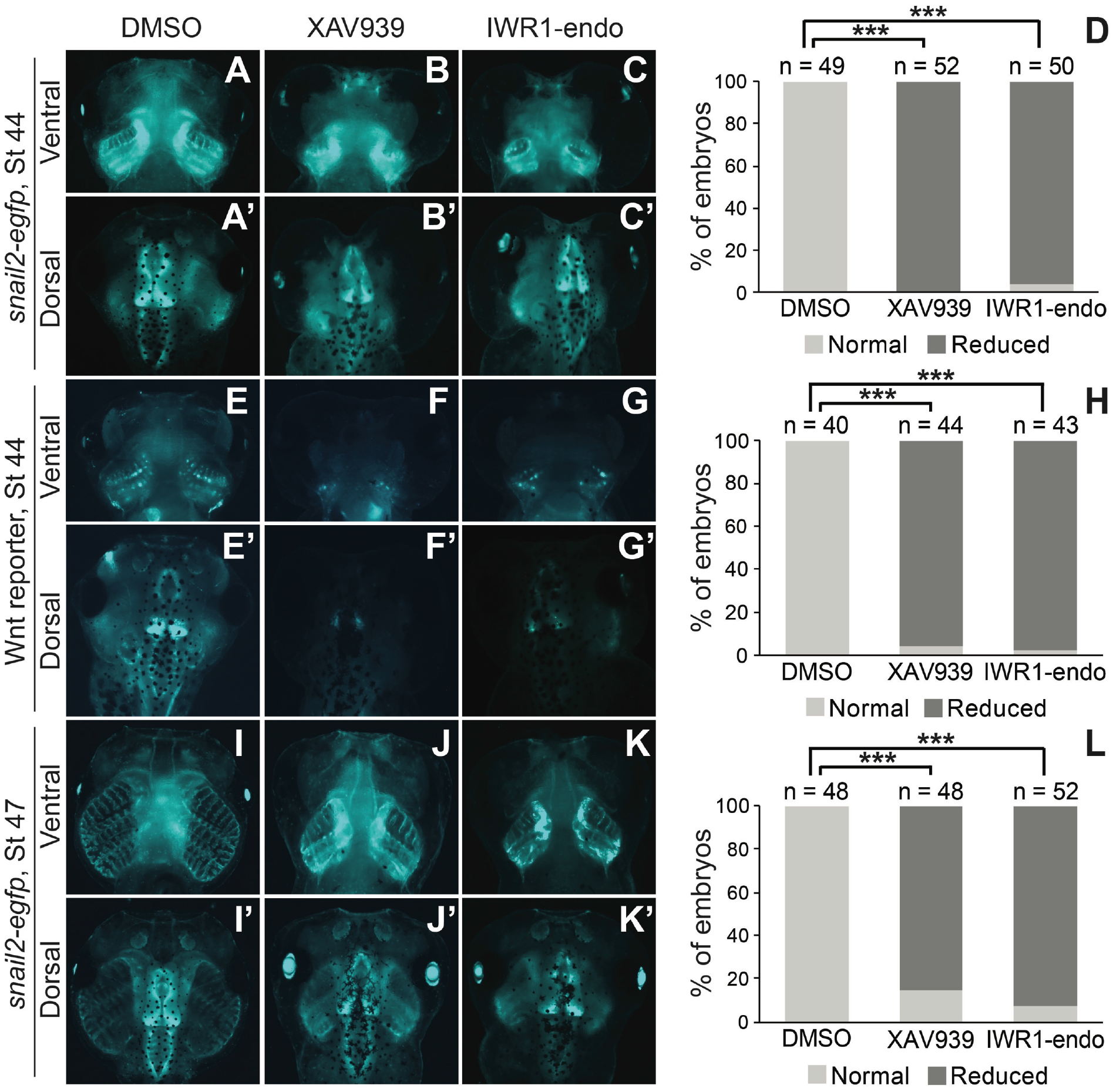
Wnt signaling is required for the differentiation of CNC into head cartilage structures. *Snail2::egfp* or Wnt reporter embryos were treated with XAV939 (**B, B’, F, F’**, 20 μM; **J** and **J’**, 5 μM), IWR1-endo (**C, C’, G, G’**, 40 μM; **K** and **K’**, 10 μM) or DMSO (vehicle control) from stage ~28 to ~35. Embryos were washed and cultured again to the indicated stages. A representative embryo from each treatment group is shown on the left, with upper and lower panels displaying ventral and dorsal views, respectively, of the same embryos, and statistics is shown in the graphs on the right. ***, P < 0.001.

Finally, we examined directly if Wnt is responsible for inducing *snail2* expression in the differentiating CNC. As shown in Fig. 7A-C, G, incubation of wild-type embryos in low-dosage XAV939 or IWR1-endo starting at stage ~28 resulted in decreased *snail2* expression at stage ~35 in the condensing mesenchyme in pharyngeal arches, suggesting that Wnt is required for *snail2* expression in the post-migratory CNC that are about to differentiate into head cartilage structures. Previous studies have shown that knockdown of Snail2 causes loss of *sox9* transcripts during *Xenopus* CNC induction^12,40^. Because Sox9 is a skeletogenic CNC marker and a master regulator of chondrogenesis from the CNC lineage at later stages^41,42^, we assessed the effects of Wnt inhibition after CNC migration on the expression of *sox9*. Similar to *snail2*, inhibition of Wnt signaling after CNC migration also reduced the expression of *sox9* at stage ~35 (Fig. 7D-F, H), providing a possible mechanism for the inhibition of head cartilage differentiation as shown in Fig. 6. Taken together, these data indicate that Wnt signaling is indispensable for *sox9* expression and head cartilage differentiation, possibly through inducing *snail2* expression.

**Figure 7.**
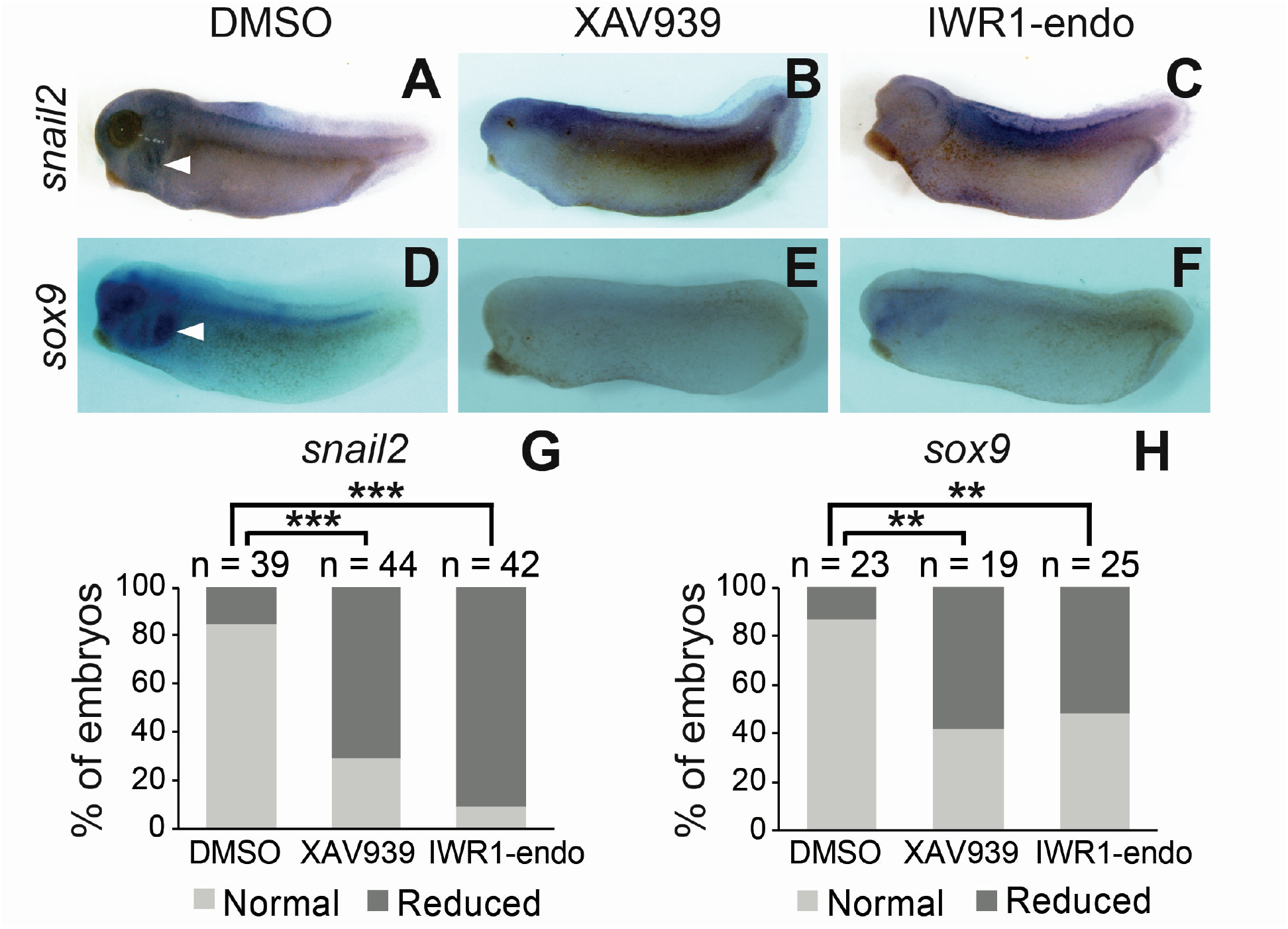
Inhibition of Wnt signaling reduces *snail2* and *sox9* expression in the post-migratory CNC. Wild-type embryos were treated with XAV939 (5 μM), IWR1-endo (10 μM) or DMSO from stage ~28 to ~35, and processed for in situ hybridization for *snail2* or *sox9*.A representative tadpole from each treatment group is shown in the upper panels (**A-C**, *snail2*; **D-F**, sox9), and statistics is shown in the graphs (**G**, *snail2*; **H**, sox9). White arrowheads point to staining in the condensing mesenchyme in the pharyngeal arches, which is reduced in tadpoles treated with Wnt inhibitors. **, P < 0.01; ***, P < 0.001.

## Discussion

The CNC can differentiate into many types of cells during early embryonic development^1–3^. Transgenic reporter animals provide powerful lineage-tracing tools for identifying CNC derivatives and understanding the mechanisms that control CNC differentiation, which are critical for the studies of CNC biology as well as the prevention and treatment of neurocristopathies. Although *X. laevis* and *X. tropicalis* have long been used to study CNC induction and migration^8^, little is known about CNC differentiation in these species. Instead, most of our knowledge on CNC differentiation was obtained from previous transgenic studies using other models such as mice and zebrafish^7–9,34^. Most recently, the first two *X. laevis* transgenic CNC reporter lines were generated. However, these lines are suitable for imaging CNC induction and migration, respectively, but not differentiation^11^. Here we report the first *X. tropicalis* transgenic CNC reporter line, which can be used not only for tracing CNC induction and early migration, but also for high-resolution live imaging of CNC differentiation. The ability of eGFP to label CNC derivatives is due to the expression of *snail2* in the differentiating CNC, which has not been described before.

Snail2 is a transcription factor that is expressed in the early CNC in vertebrates^26,27^, but the timing of *snail2* expression in the CNC varies in different species. Transcripts of *snail2* are detectable in the pre-migratory CNC in frogs, reptiles and chicks, but not in fish or mice^26,27,43^. Snail2 in the pre-migratory CNC is required for the emigration of CNC cells from the neural tube in frogs and chicks, likely due to its ability to induce EMT^16,43^. At later stages, *snail2* transcripts were found in the migrating CNC in essentially all vertebrate species that have been examined, including *Xenopus*, chicks and mice^13,27,43^. It has also been shown that *snail2* expression diminishes toward the end of CNC migration in both *Xenopus* and chick embryos^17–19,43^. This is in line with the hypothesis that CNC cells undergo mesenchymal-to-epithelial transition, which is the reciprocal of EMT, to stop migration, allowing them to colonize various tissues in the embryo^1^. Currently, there is no published information on the expression of *snail2* after CNC migration in any species. However, studies with mice suggest that Snail2 may function in CNC development at later stages. While whole-embryo double knockout of *snail2* and its close paralog *snail1* has no effect on CNC induction or early migration^44^, neural crest-specific loss of *snail1* on the *snail2*-null background leads to multiple craniofacial defects that are reminiscent of conditional neural crest mutants of several other important genes^45^. In contrast, neither conditional knockout of *snail1* in the neural crest nor global knockout of *snail2* alone causes these defects^45^. These results imply a redundant role of Snail1 and Snail2 in late CNC development, possibly in CNC differentiation. Therefore, it is important to determine if the Snail family of transcription factors are expressed and functional in the post-migratory CNC. Here we report that *snail2* is re-expressed in the differentiating CNC in *X. tropicalis* embryos. This is supported by the fluorescence pattern displayed by the *snail2::egfp* transgenic tadpoles, as well as the in situ hybridization and immunohistochemistry data. Our results are consistent with a possible role of *snail2* during CNC differentiation, as implicated by the mouse study^45^. It remains to be examined if *snail2* is similarly expressed in the differentiating CNC in mice and other vertebrates, and if this gene indeed functions in CNC differentiation.

The Wnt signaling pathway is a major inducer of Snail2 in various vertebrate species, mainly through direct activation of *snail2* transcription^15,46^. In addition, Wnt-induced GSK3β inhibition can lead to stabilization of the Snail2 protein^47^, which is capable of binding to its own enhancer and further stimulating *snail2* expression^48,49^. In *Xenopus* embryos, Wnt signaling induces the formation of the NPB. After NPB formation, a second wave of Wnt signal activates the expression of Snail2, which is required for CNC specification within the NPB^50^. Both the Wnt signal and Snail2 mRNA/protein are clearly detectable throughout the pre-migratory CNC (Fig. 2B-D, 3A-A”, 5A, B). The accumulating Snail2 in the pre-migratory CNC likely prepares the CNC cells for EMT/migration, as knockdown of Snail2 inhibits CNC migration^16^. It has also been shown that in pre-migratory *Xenopus* CNC explants, β-catenin is mainly detected in the nucleus; in contrast, in migrating CNC explants, β-catenin re-localizes to the plasma membrane, indicating a reduction of Wnt signaling in CNC cells that have emigrated from the neural tube^51^. Because Wnt induces *snail2* expression, these observations provide a possible mechanism for the downregulation of *snail2* transcripts during CNC migration. Our data further suggest that the reexpression of *snail2* during CNC differentiation is also driven by Wnt signaling. A comparison between the fluorescence patterns of *snail2::egfp* and Wnt reporter transgenic tadpoles shows striking similarity during CNC differentiation, and blocking Wnt signaling after CNC migration inhibits *snail2* expression and head cartilage differentiation (Fig. 1D-E”, 5D-E’, 6, 7). Thus, the Wnt-Snail2 axis may function reiteratively during CNC specification, emigration and differentiation.

Wnt signaling is known to be important for neural crest differentiation, but the exact roles of Wnt in this developmental process are controversial and may vary from species to species. An earlier report shows that Wnt promotes CNC differentiation into pigment cells at the expense of neurons and glia in zebrafish^52^. In contrast, β-catenin instructs mouse neural crest cells to adopt a sensory neuronal fate at the cost of essentially all other neural crest derivatives, presumably through mediating Wnt signaling^53,54^. Thus, the effects of Wnt signaling on cell fate determination during neural crest differentiation seem to be species-dependent. With regard to craniofacial morphogenesis, Wnt is crucial to the selection between chondrocytic and osteoblastic fates in the mammalian CNC. Specifically, Wnt promotes bone formation and simultaneously suppresses chondrogenesis in mice^34^. However, Wnt has also been shown to be required for chondrogenic differentiation in cultured mouse cells in a Sox9-dependent manner^55^. In zebrafish, blocking Wnt signaling after CNC migration inhibits ventral cartilage differentiation^56^. Although head cartilage is often used as a phenotypic readout for disrupted CNC induction or migration in *Xenopus*, little is known about how CNC cells differentiate into cartilaginous structures in frogs. This is probably because genes and pathways that are important for CNC differentiation, such as Wnt and *sox9*, often play critical roles in earlier induction and/or migration as well, and tools for temporally controlled gene inactivation in *Xenopus* are lacking. In the current study, we show that treating *X. tropicalis* embryos with Wnt inhibitors after CNC migration leads to reduced *sox9* expression and under-differentiated head cartilage structures (Fig. 6, 7). To our knowledge, this is the first evidence of Wnt function in *Xenopus* CNC differentiation. Future studies are needed to understand how Wnt affects *sox9* expression and head cartilage differentiation in *Xenopus*.

The *snail2::egfp* transgenic line is a useful tool for live imaging of CNC development in *Xenopus*. Normal and dysregulated CNC induction, migration and differentiation can be visualized directly in the transgenic embryos (Fig. 1, 4), making them highly suitable for high-throughput screens to identify genetic and environmental factors that interfere with CNC development at any stage. The labeling of multiple CNC derivatives, such as cells in the head cartilage, cranial nerves, and thymus, by eGFP in the *snail2::egfp* transgenic tadpoles (Fig. 1E-H’), further raises the possibility of using this transgenic line to identify new types of cells that derive from the CNC. For example, there is published evidence supporting a CNC origin for the gonadotropin releasing hormone positive and microvillous neurons in the early zebrafish olfactory epithelium, but a most recent study suggests that all the sensory neurons in the zebrafish olfactory epithelium derive from the preplacodal ectoderm instead^22,57,58^. Interestingly, we observed clear eGFP expression in the olfactory epithelium of *snail2::egfp* tadpoles starting at stage ~42, when the microvillous neurons just emerge^59^. Whether there is a CNC contribution to the *Xenopus* olfactory epithelium warrants further investigation, and additional transgenic tools may be needed for this type of studies. To better trace the dynamic development of CNC cells, we are in the process of generating a new *snail2::mEos3.2* line, in which the CNC lineage is labeled with the mEOS3.2 photoswitching fluorescent protein. This second-generation *snail2* reporter line will allow the labeling of pre-migratory or migrating CNC cells and tracing their fates at later stages. Together, these transgenic reporter lines should have a profound impact on the studies of CNC development.

## Methods

### Plasmids and antibodies

Genomic DNA was prepared from *X. tropicalis* embryos as described^60^, and a 3.9 kb fragment of the *snail2* promoter/enhancer, as reported by Vallin et al.^15^, was cloned using nested PCR. To generate the transgenic construct, the *snail2* promoter/enhancer sequence was subcloned into the IS-egfp transgenic vector (a gift from Dr. Robert Grainger)^20,21^. Primers used in cloning and subcloning are listed in Table S1. Constructs for preparing the in situ hybridization probes for *snail2* and *sox9* were obtained previously^28^. The mouse anti-Snail2 (DSHB 62.1E6, 1:50) and rabbit anti-GFP (Life Technologies A11122, 1:200), as well as Alexa Fluor 594 AffiniPure donkey anti-mouse and Alexa Fluor 488 AffiniPure donkey anti-rabbit antibodies (Jackson Immuno Research Laboratories 715-585-150 and 711-545-152, 1:500 for both), were used for immunohistochemistry.

### Animals and transgenesis

Wild-type *X. tropicalis* adults (male and female) were purchased from NASCO, and the Wnt reporter line was generated in a previous study^32^. ISceI-mediated transgenesis was carried out as described by Ogino et al. to generate the *snail2::egfp* transgenic founders^20^. Briefly, the IS-*snail2::egfp* plasmid was digested with I-SceI enzyme, and the reaction mixture was injected into fertilized *X. tropicalis* eggs. Embryos with potential non-mosaic eGFP expression were selected and raised to adulthood. These transgenic founders were crossed with wild-type *X. tropicalis* frogs to generate heterozygotes, which were further inbred to obtain homozygotes.

### Embryo manipulation

Embryo were obtained by natural mating and cultured in 0.1x MBS to desired stages as described previously^28^. MO 13-3, the antisense MO for ADAM13, was synthesized by Gene Tools, and the sequence was reported previously^28^. For injections, 8-cell stage *snail2::egfp* embryos were injected in a single dorsal-animal blastomere with 1.5 ng MO 13-3 using a PLI-100A microinjector (Harvard Apparatus), and Alexa Fluor 555 dextran (Invitrogen) was coinjected as a lineage tracer. For in situ hybridization and immunohistochemistry, embryos were fixed at desired stages and processed as described^60^. For Wnt inhibitor treatment, embryos were cultured in XAV939 or IWR1-endo (both were from Selleckchem) from stage ~28 to ~35. *Snail2-egfp* or Wnt reporter transgenic tadpoles were washed three times and subsequently cultured in 0.1x MBS until stage ~44 or ~47 (Fig. 6); wild-type tadpoles were immediately fixed and processed for in situ hybridization for *snail2* or *sox9* (Fig. 7). Fluorescent and bright-field images were taken with a Zeiss Axiozoom.V16 epifluorescence microscope. Image acquisition and processing were carried out using an AxioCam MRc Rev3 camera and the ZEN 2.0 software package.

### Phenotype scoring and statistics

Injected embryos were scored by comparing the injected side with the uninjected side of the same embryos, and Wnt inhibitor-treated embryos were scored by comparing with the DMSO-treated controls. The percentage of normal and reduced phenotypes were calculated, and Chi-squared tests were performed to compare the phenotypes in different treatment groups.

### Ethics statement

Methods involving live animals were carried out in accordance with the guidelines and regulations approved and enforced by the Institutional Animal Care and Use Committee at West Virginia University and the University of Delaware.

## Acknowledgements

We thank Drs. Robert Grainger, Takuya Nakayama and Anoop Shah from the University of Virginia for help with transgenesis. This work was funded by the U.S. NIH (R01GM114105, R03DE022813 and P20GM104316 to SW, and R01EY015279 to MKD) and March of Dimes Foundation (1-FY10-399 to SW). Additionally, part of the imaging work was supported by NIH grant 1S10RR027273-01. Research in the Vleminckx laboratory is supported by the Research Foundation – Flanders (FWO-Vlaanderen) (grants G0A1515N and G029413N) and by the Concerted Research Actions from Ghent University (B0F15/G0A/011); further support was obtained from the Hercules Foundation, Flanders (grant AUGE/11/14).

## Author contributions

JL, MP, MKD and SW designed the experiments. JL, MP, CM, RL and SW performed the experiments; HTT and KV generated the transgenic Wnt reporter line and provided assistance on experiments using this transgenic line. JL, MP and SW analyzed the data and wrote the manuscript; MKD and KV provided comments and assistance on manuscript preparation.

## Data availability

All data generated or analysed during this study are included in this published article (and its Supplementary Information files).

## References

1. Simões-Costa, M. & Bronner, M.E. Establishing neural crest identity: a gene regulatory recipe. Development 142, 242–57 (2015).

2. Bronner, M.E. & Simões-Costa, M. The Neural Crest Migrating into the Twenty-First Century. Curr Top Dev Biol 116, 115–34 (2016).

3. Mayor, R. & Theveneau, E. The neural crest. Development 140, 2247–51 (2013).

4. Barraud, P. et al. Neural crest origin of olfactory ensheathing glia. Proc Natl Acad Sci U S A 107, 21040–5 (2010).

5. Mongera, A. et al. Genetic lineage labeling in zebrafish uncovers novel neural crest contributions to the head, including gill pillar cells. Development 140, 916–25 (2013).

6. Trainor, P.A. Developmental Biology: We Are All Walking Mutants. Curr Top Dev Biol 117, 523–38 (2016).

7. Aoto, K. et al. Mef2c-F10N enhancer driven β-galactosidase (LacZ) and Cre recombinase mice facilitate analyses of gene function and lineage fate in neural crest cells. Dev Biol 402, 3–16 (2015).

8. Barriga, E.H., Trainor, P.A., Bronner, M. & Mayor, R. Animal models for studying neural crest development: is the mouse different? Development 142, 1555–60 (2015).

9. Carney, T.J. et al. A direct role for Sox10 in specification of neural crest-derived sensory neurons. Development 133, 4619–30 (2006).

10. Rodrigues, F.S., Doughton, G., Yang, B. & Kelsh, R.N. A novel transgenic line using the Cre-lox system to allow permanent lineage-labeling of the zebrafish neural crest. Genesis 50, 750–7 (2012).

11. Alkobtawi, M. et al. Characterization of Pax3 and Sox10 transgenic Xenopus laevis embryos as tools to study neural crest development. Dev Biol (2018).

12. Shi, J., Severson, C., Yang, J., Wedlich, D. & Klymkowsky, M.W. Snail2 controls mesodermal BMP/Wnt induction of neural crest. Development 138, 3135–45 (2011).

13. Mayor, R., Morgan, R. & Sargent, M.G. Induction of the prospective neural crest of Xenopus. Development 121, 767–77 (1995).

14. Wu, J., Saint-Jeannet, J.P. & Klein, P.S. Wnt-frizzled signaling in neural crest formation. Trends Neurosci 26, 40–5 (2003).

15. Vallin, J. et al. Cloning and characterization of three Xenopus slug promoters reveal direct regulation by Lef/beta-catenin signaling. J Biol Chem 276, 30350–8 (2001).

16. Carl, T.F., Dufton, C., Hanken, J. & Klymkowsky, M.W. Inhibition of neural crest migration in Xenopus using antisense slug RNA. Dev Biol 213, 101–15 (1999).

17. Huang, C., Kratzer, M.C., Wedlich, D. & Kashef, J. E-cadherin is required for cranial neural crest migration in Xenopus laevis. Dev Biol 411, 159–71 (2016).

18. Owens, N.D. et al. Measuring Absolute RNA Copy Numbers at High Temporal Resolution Reveals Transcriptome Kinetics in Development. Cell Rep 14, 632–47 (2016).

19. Session, A.M. et al. Genome evolution in the allotetraploid frog Xenopus laevis. Nature 538, 336–343 (2016).

20. Ogino, H., McConnell, W.B. & Grainger, R.M. High-throughput transgenesis in Xenopus using I-SceI meganuclease. Nat Protoc 1, 1703–10 (2006).

21. Ogino, H., McConnell, W.B. & Grainger, R.M. Highly efficient transgenesis in Xenopus tropicalis using I-SceI meganuclease. Mech Dev 123, 103–13 (2006).

22. Saxena, A., Peng, B.N. & Bronner, M.E. Sox10-dependent neural crest origin of olfactory microvillous neurons in zebrafish. Elife (Cambridge) 2, e00336 (2013).

23. Suzuki, J., Yoshizaki, K., Kobayashi, T. & Osumi, N. Neural crest-derived horizontal basal cells as tissue stem cells in the adult olfactory epithelium. Neurosci Res 75, 112–20 (2013).

24. Lee, Y.H. et al. Early development of the thymus in Xenopus laevis. Dev Dyn 242, 164–78 (2013).

25. Corish, P. & Tyler-Smith, C. Attenuation of green fluorescent protein half-life in mammalian cells. Protein Eng 12, 1035–40 (1999).

26. Locascio, A., Manzanares, M., Blanco, M.J. & Nieto, M.A. Modularity and reshuffling of Snail and Slug expression during vertebrate evolution. Proc Natl Acad Sci U S A 99, 16841–6 (2002).

27. Jiang, R., Lan, Y., Norton, C.R., Sundberg, J.P. & Gridley, T. The Slug gene is not essential for mesoderm or neural crest development in mice. Dev Biol 198, 277–85 (1998).

28. Wei, S. et al. ADAM13 induces cranial neural crest by cleaving class B Ephrins and regulating Wnt signaling. Dev Cell 19, 345–52 (2010).

29. Cousin, H., Abbruzzese, G., Kerdavid, E., Gaultier, A. & Alfandari, D. Translocation of the cytoplasmic domain of ADAM13 to the nucleus is essential for Calpain8-a expression and cranial neural crest cell migration. Dev Cell 20, 256–63 (2011).

30. Li, J. et al. ADAM19 regulates Wnt signaling and neural crest specification by stabilizing ADAM13. Development 145(2018).

31. Wei, S. et al. Roles of ADAM13-regulated Wnt activity in early Xenopus eye development. Dev Biol 363, 147–54 (2012).

32. Tran, H.T., Sekkali, B., Van Imschoot, G., Janssens, S. & Vleminckx, K. Wnt/beta-catenin signaling is involved in the induction and maintenance of primitive hematopoiesis in the vertebrate embryo. Proc Natl Acad Sci U S A 107, 16160–5 (2010).

33. Borday, C. et al. An atlas of Wnt activity during embryogenesis in Xenopus tropicalis. PLoS One 13, e0193606 (2018).

34. Bhatt, S., Diaz, R. & Trainor, P.A. Signals and switches in Mammalian neural crest cell differentiation. Cold Spring Harb Perspect Biol 5(2013).

35. Kurosaka, H., Trainor, P.A., Leroux-Berger, M. & Iulianella, A. Cranial nerve development requires co-ordinated Shh and canonical Wnt signaling. PLoS One 10, e0120821 (2015).

36. Wang, Y. et al. Spatiotemporal dynamics of canonical Wnt signaling during embryonic eye development and posterior capsular opacification (PCO). Exp Eye Res 175, 148–158 (2018).

37. Noelanders, R. & Vleminckx, K. How Wnt Signaling Builds the Brain: Bridging Development and Disease. Neuroscientist (2016).

38. Huang, S.M. et al. Tankyrase inhibition stabilizes axin and antagonizes Wnt signalling. Nature 461, 614–20 (2009).

39. Chen, B. et al. Small molecule-mediated disruption of Wnt-dependent signaling in tissue regeneration and cancer. Nat Chem Biol 5, 100–7 (2009).

40. Zhang, C., Carl, T.F., Trudeau, E.D., Simmet, T. & Klymkowsky, M.W. An NF-kappaB and slug regulatory loop active in early vertebrate mesoderm. PLoS One 1, e106 (2006).

41. Mori-Akiyama, Y., Akiyama, H., Rowitch, D.H. & de Crombrugghe, B. Sox9 is required for determination of the chondrogenic cell lineage in the cranial neural crest. Proc Natl Acad Sci U S A 100, 9360–5 (2003).

42. Lee, Y.H. & Saint-Jeannet, J.P. Sox9 function in craniofacial development and disease. Genesis 49, 200–8 (2011).

43. Nieto, M.A., Sargent, M.G., Wilkinson, D.G. & Cooke, J. Control of cell behavior during vertebrate development by Slug, a zinc finger gene. Science 264, 835–9 (1994).

44. Murray, S.A. & Gridley, T. Snail family genes are required for left-right asymmetry determination, but not neural crest formation, in mice. Proc Natl Acad Sci U S A 103, 10300–4 (2006).

45. Murray, S.A., Oram, K.F. & Gridley, T. Multiple functions of Snail family genes during palate development in mice. Development 134, 1789–97 (2007).

46. Taneyhill, L.A. & Bronner-Fraser, M. Dynamic alterations in gene expression after Wnt-mediated induction of avian neural crest. Mol Biol Cell 16, 5283–93 (2005).

47. Wu, Z.Q. et al. Canonical Wnt signaling regulates Slug activity and links epithelial-mesenchymal transition with epigenetic Breast Cancer 1, Early Onset (BRCA1) repression. Proc Natl Acad Sci U S A 109, 16654–9 (2012).

48. Sakai, D., Suzuki, T., Osumi, N. & Wakamatsu, Y. Cooperative action of Sox9, Snail2 and PKA signaling in early neural crest development. Development 133, 1323–33 (2006).

49. LaBonne, C. & Bronner-Fraser, M. Snail-related transcriptional repressors are required in Xenopus for both the induction of the neural crest and its subsequent migration. Dev Biol 221, 195–205 (2000).

50. Stuhlmiller, T.J. & Garcia-Castro, M.I. Current perspectives of the signaling pathways directing neural crest induction. Cell Mol Life Sci 69, 3715–37 (2012).

51. Maj, E. et al. Controlled levels of canonical Wnt signaling are required for neural crest migration. Dev Biol 417, 77–90 (2016).

52. Dorsky, R.I., Moon, R.T. & Raible, D.W. Control of neural crest cell fate by the Wnt signalling pathway. Nature 396, 370–3 (1998).

53. Lee, H.Y. et al. Instructive role of Wnt/beta-catenin in sensory fate specification in neural crest stem cells. Science 303, 1020–3 (2004).

54. Hari, L. et al. Lineage-specific requirements of beta-catenin in neural crest development. J Cell Biol 159, 867–80 (2002).

55. Yano, F. et al. The canonical Wnt signaling pathway promotes chondrocyte differentiation in a Sox9-dependent manner. Biochem Biophys Res Commun 333, 1300–8 (2005).

56. Alexander, C., Piloto, S., Le Pabic, P. & Schilling, T.F. Wnt signaling interacts with bmp and edn1 to regulate dorsal-ventral patterning and growth of the craniofacial skeleton. PLoS Genet 10, e1004479 (2014).

57. Whitlock, K.E., Smith, K.M., Kim, H. & Harden, M.V. A role for foxd3 and sox10 in the differentiation of gonadotropin-releasing hormone (GnRH) cells in the zebrafish Danio rerio. Development 132, 5491–502 (2005).

58. Aguillon, R. et al. Cell-type heterogeneity in the early zebrafish olfactory epithelium is generated from progenitors within preplacodal ectoderm. Elife 7(2018).

59. Hansen, A., Reiss, J.O., Gentry, C.L. & Burd, G.D. Ultrastructure of the olfactory organ in the clawed frog, Xenopus laevis, during larval development and metamorphosis. J Comp Neurol 398, 273–88 (1998).

60. Sive, H.L., Grainger, R.M. & Harland, R.M. Early Development of Xenopus Laevis. A Laboratory Manual. Cold Spring Harbor Laboratory Press. (2000).

