## Supplemental information for "A new transgenic reporter line reveals Wnt-dependent Snail2 reexpression and cranial neural crest differentiation in *Xenopus*"

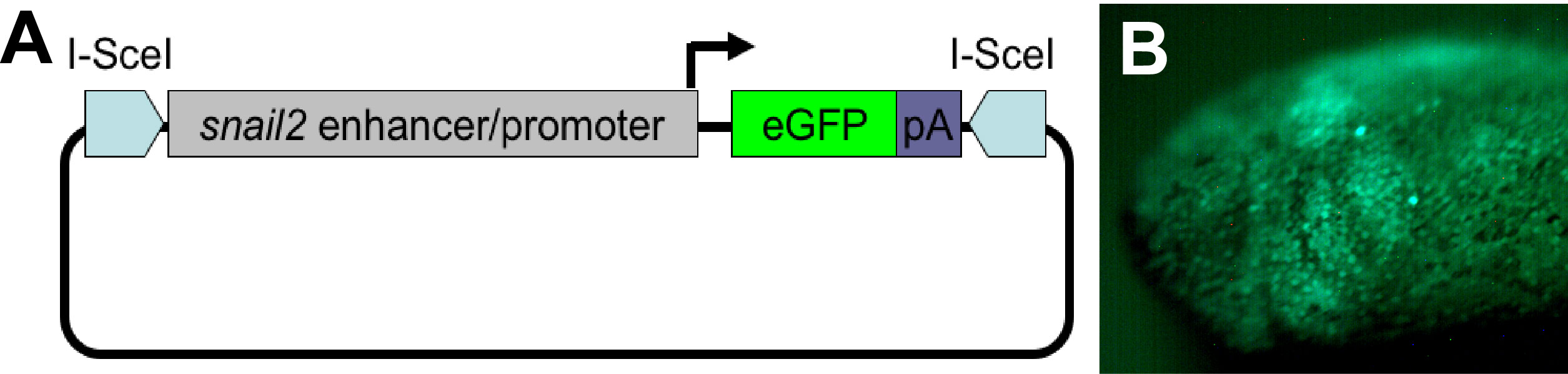

**Supplemental Fig. S1. Generation of the *snail2::egfp* transgenic line. A.** The transgenic construct that was used for generating the *snail2::egfp* line. pA, SV40 polyadenylation site. **B.** A *snail2::egfp* transgenic founder showing eGFP expression in the migrating CNC streams.

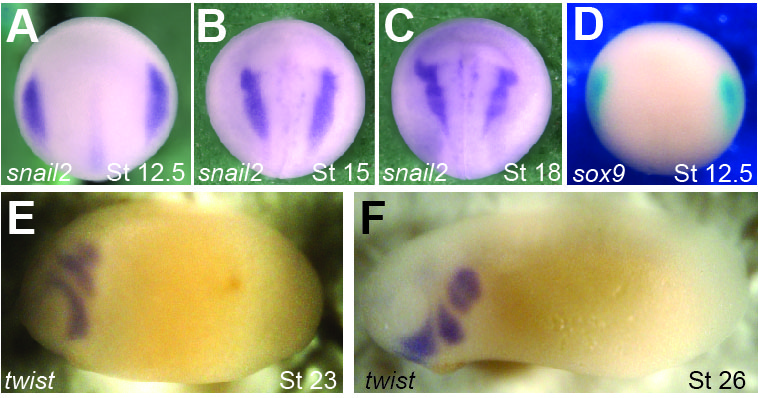

**Supplemental Fig. S2. Normal marker expression in the pre-migratory and migrating CNC in heterozygous *snail2::egfp* transgenic embryos.** Heterozygous *snail2::egfp* embryos were fixed at the indicated stages, and in situ hybridization was carried out for *snail2* (**A**-**C**), sox9 (**D**) or *twist* (**E** and **F**).

**Table S1 Primers used for cloning and subcloning of the *snail2* promoter/enhancer**

| Primer name | Sequence (5’-3’) |
| --- | --- |
| *Snail2* forward | GGCCCTGTACATTGTTGG |
| *Snail2* reverse | AGAGGCACAGGATCTGCATTACTG |
| IS-*snail2*::eGFP SalI forward | ATATGTCGACGGCCCTGTACATTGTTGG |
| IS-*snail2*::eGFP NotI reverse | ATATGCGGCCGCTCACCAAGTGGCAGCGCT |

Restriction sites that were used to subclone the *snail2* promoter/enhancer into the IS-eGFP vector are underscored.
